# Non-invasive homogeneous targeted blood-brain barrier disruption using acoustic holography with a clinical focused ultrasound system

**DOI:** 10.1101/2023.12.05.570091

**Authors:** Nathan McDannold, Yongzhi Zhang, Stecia-Marie Fletcher, Margaret Livingstone

**Affiliations:** Department of Radiology, Brigham and Women’s Hospital, Harvard Medical School; Department of Neurobiology, Harvard Medical School

## Abstract

Holographic methods can be used with phased array transducers to shape an ultrasound field. We tested a simple method to create holograms with a 1024-element phased array transducer. With this method, individual acoustic simulations for each element of the transducer were simultaneously loaded into computer memory. Each element’s phase was systematically modulated until the combined field matched a desired pattern. The method was evaluated with a 220 kHz hemispherical transducer being tested clinically to enhance drug delivery via blood-brain barrier disruption. The holograms were evaluated in a tissue-mimicking phantom and *in vivo* in experiments disrupting the blood-brain barrier in rats and in a macaque. This approach can enlarge the focal volume in a patient-specific manner and could reduce the number of sonication targets needed to disrupt large volumes, improve the homogeneity of the disruption, and improve our ability to detect microbubble activity in tissues with low vascular density.

**Teaser:** Holography can shape the focal region of a clinical focused ultrasound system developed for targeted drug delivery in the brain.

## Introduction

As with light, holography can be used with ultrasound to create complex acoustic patterns using phased arrays transducers or acoustic lenses (*1-4*). Acoustic holograms have been used to tailor the sound field to a desired shape for various applications, such as acoustic tweezers, ultrasound ablation, neuromodulation, and tactile displays.

One promising therapeutic ultrasound application uses microbubble-enhanced focused ultrasound to temporarily disrupt the blood-brain barrier (BBB) (*5*) and target the delivery of drugs that normally do not extravasate in the brain. This technique is currently being tested in several clinical trials (*6-9*). For many applications, large volume – even whole brain – BBB disruption is desired, which requires ultrasound exposures (sonications) at hundreds of individual targets. While a phased array can do this rapidly, it would be beneficial to enlarge the focal region and tailor its shape to match different anatomical structures, enable more uniform disruption, and reduce the overall number of sonication targets. Such an approach might also aid in controlling the procedure, which is currently done by monitoring the acoustic emissions emitted by the oscillating microbubbles in the ultrasound field (*10*). Signals arising from sonication in tissues with low vascular density, such as white matter tracts, are weak and can be challenging to monitor during transcranial sonication (*11*). Enlarging and shaping the focal volume could tailor the exposures to homogenous tissue structures while at the same time increasing the strength of the acoustic emissions.

Here we tested a simple method to generate acoustic holograms for phased arrays with arbitrary geometries. The method was evaluated for a clinical 1024-element phased array system currently being tested clinically for BBB disruption (*6-9*). The holograms were verified using MR temperature imaging (*12*) and tested for enlarging the focal region in rats and a macaque monkey.

## Results

Holograms were created by loading 1024 individual acoustic simulations for the elements of the ExAblate Neuro Type II phased array transducer (InSightec) simultaneously into memory and systematically modulating each element’s phase to maximize agreement between the three-dimensional pressure distribution and a desired goal (Fig. 1). To reduce memory requirements, the simulations covered only a small volume (dimensions: 9.7-19.4 mm) around the geometric focus. Using a desktop PC and MATLAB, this procedure required approximately 20 minutes per hologram.

**Fig. 1.**
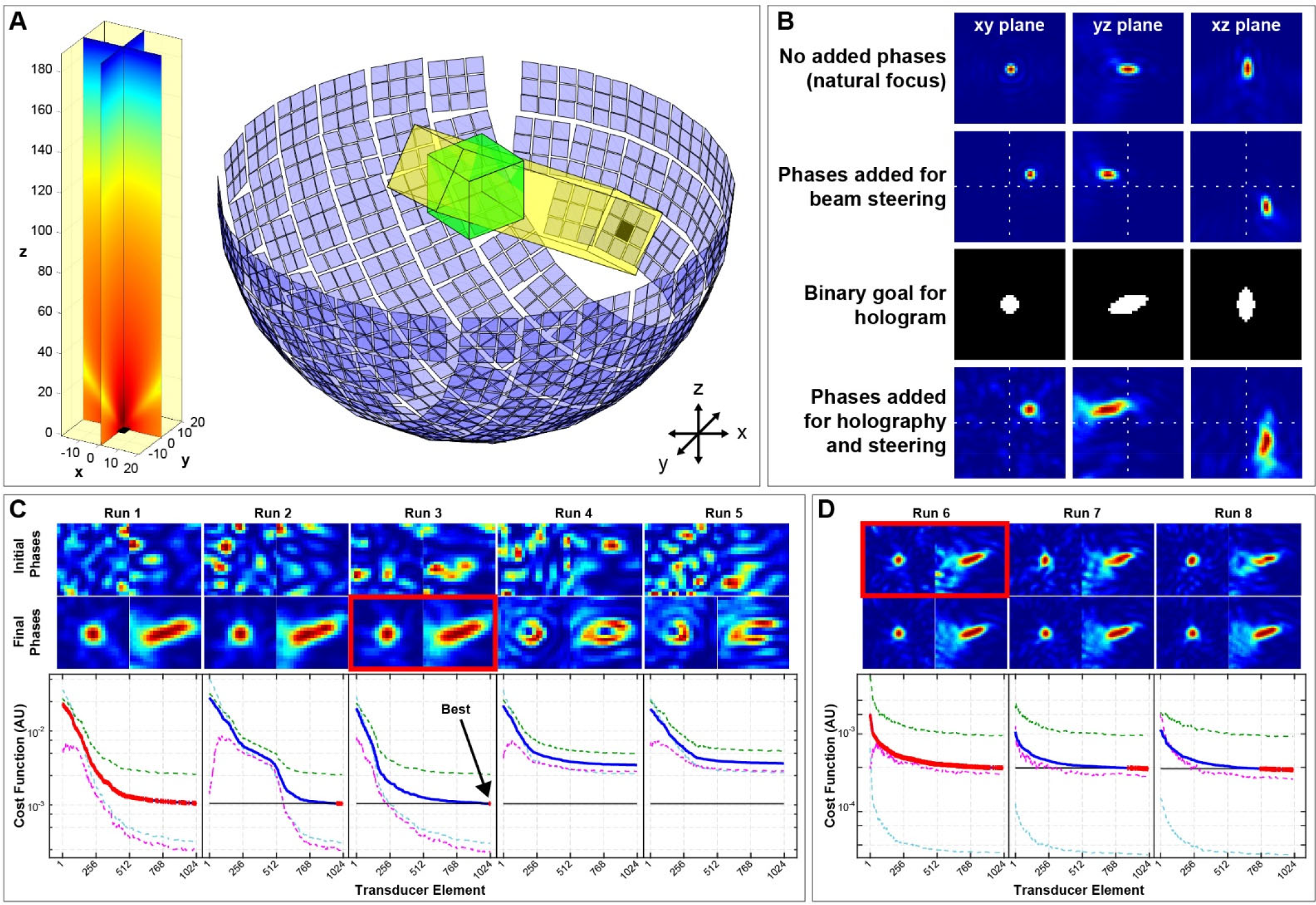
A method for holography. (**A)** The three-dimensional pressure distribution (left) for one element of the ExAblate Neuro 1024-element hemispherical phased array transducer (right) was simulated in a small volume. This array has a diameter of 30 cm and operates at 220 kHz. The acoustic field was copied, rotated, and translated to the location of each element (yellow volume), and interpolated to a global region of interest (green volume). The 74×74×69 mm^3^ simulated field for the 1024 elements could then be loaded into memory simultaneously. **(B)** A complex sum across elements produced the combined simulated field. Phase offsets can be added before summation to steer the focal point and for holography. **(C)** We cycled the phase of each element in sequence and kept those that improved agreement with our goal. Images show the initial and final fields after each run; plots show the cost function used to assess this agreement. This procedure was performed for all elements five times. Blue lines show the cost function for each run; red lines indicate when the agreement improved. The dotted lines indicate the three terms in eq. 1 used to calculate the cost function. In this example, the best solution was found in the third run; note the poor convergence in runs four and five. **(D)** Using the best solution from the prior runs as a starting point, we then repeated this procedure with a larger volume. Finally, to refine the solution, we added random phases between ± 2π/3 radians and cycled through the elements two more times.

Fig. 2 shows MR temperature imaging for various holograms obtained during sonication in a tissue-mimicking phantom. The images on the left show the binary goal mask, and the images in the center show the simulated intensity of the holographic focusing. The images on the right are MR temperature images at the end of sonications with the different holograms. The method accurately recreated the holograms. In some cases, the simulated convergence to the goal was incomplete (arrowheads in Fig. 2), and these deviations in the pressure field were evident in the thermometry.

**Fig. 2.**
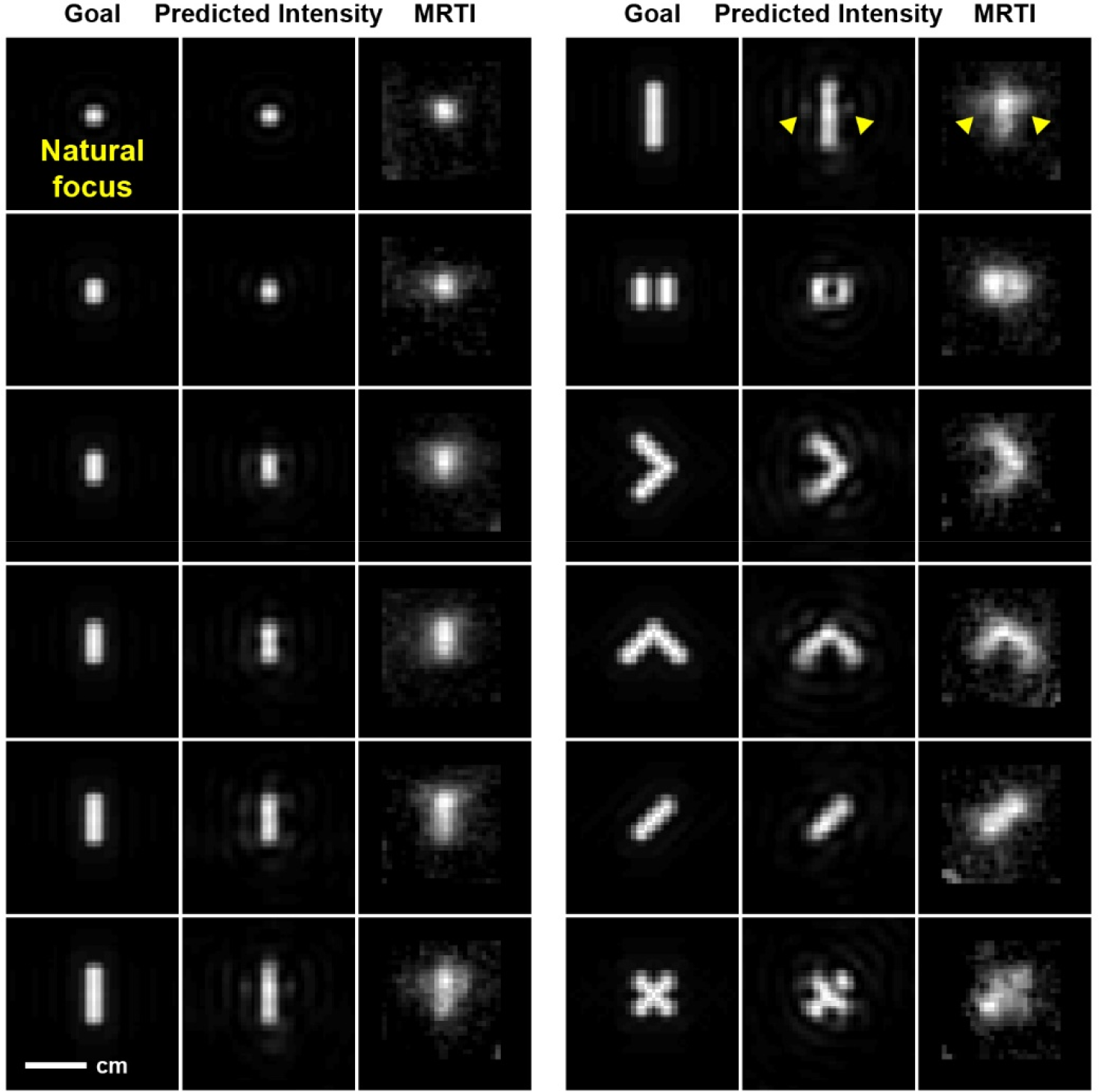
MR Temperature imaging showing heating with holographic focusing with various shapes. The left images show the binary goal, the center images show the simulated intensity, and the right images show heating. The natural focus is shown as a reference. In several cases, the holograms did not converge perfectly, and small hotspots evident in the simulations outside the binary goal were also evident in the heating (arrowheads). Images are shown in the focal (*xy*) plane of the transducer.

Next, we evaluated the method in BBB disruption experiments in rats. We first tested an elliptical focus with dimensions of 2×3×5 mm. The simulations predicted the volume of the 50% isointensity contours was 4.6 times larger than the natural focus (117.2 vs. 25.6 mm^3^). After verifying in MR temperature images that the focus shape matched the target (Fig. 3A), we applied the enlarged sonications in one hemisphere and used the natural focus in the other. The resulting BBB disruption in contrast-enhanced MRI matched the shape of the enlarged focus in the focal plane in (Fig. 3B) and covered the full thickness of the brain along the direction of the ultrasound beam. We next generated four different holograms tailored to different structures (Fig. 3C). The volumes of the 50% isointensity contours for these foci were 3.1, 3.1, 1.5, and 2.2 times that of the natural focus for locations 1-4 in Fig. 3C, respectively. The resulting contrast extravasation matched the predicted holograms reasonably well, but reflections from the back of the skull and laterally resulted in BBB disruption outside the expected volume (*).

**Fig. 3.**
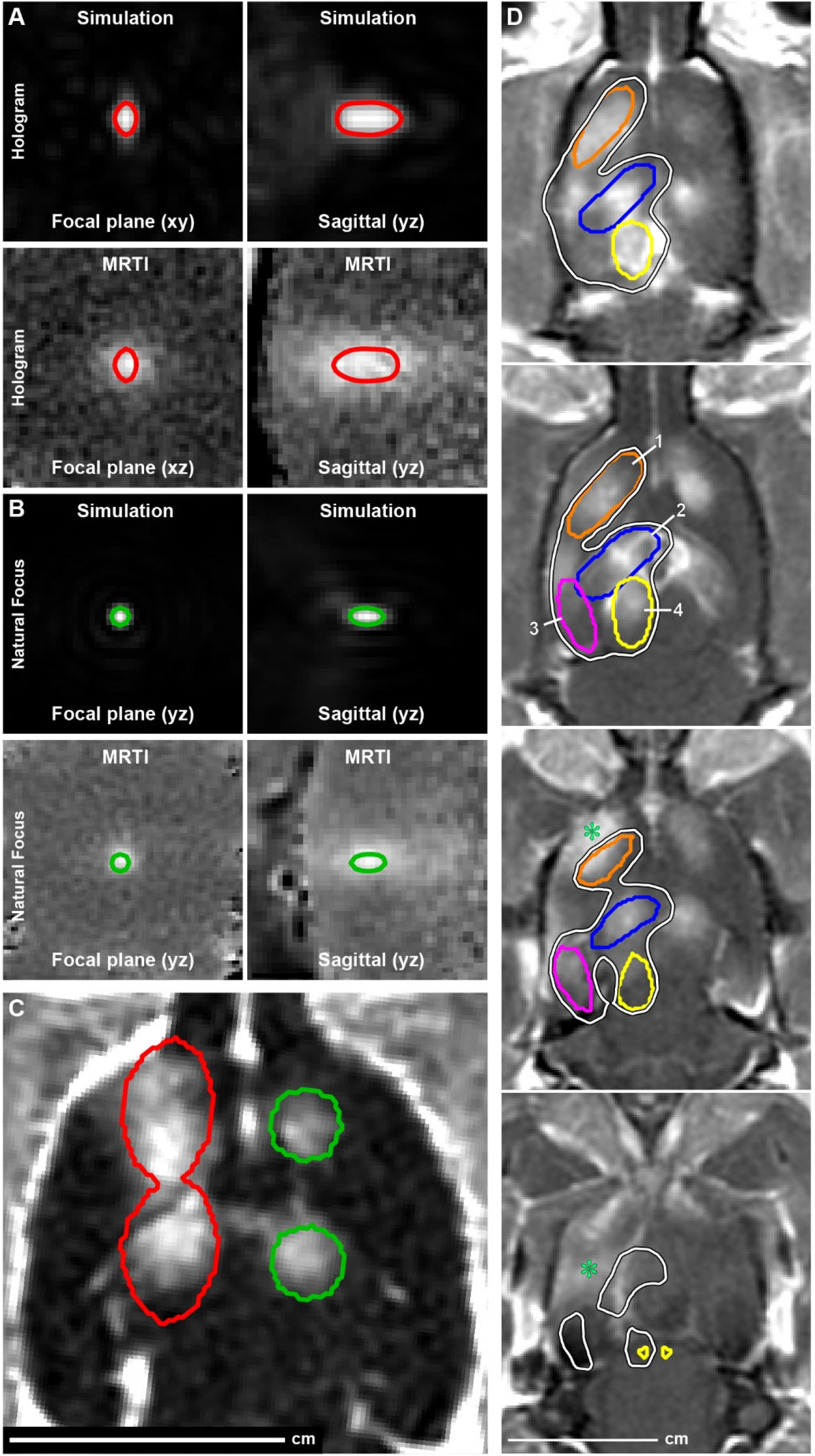
Holographic enlargement of the focus during BBB disruption in rats. **(A)** Simulated intensity maps and MR temperature imaging (MRTI) of a holographically-enlarged focus with dimensions of 2×3×5 mm. **(B)** Corresponding results using the natural focus. **(C)** Axial map (*xy* plane of transducer) showing contrast enhancement in a rat after a sonication that used the enlarged focus at two targets in one hemisphere and the natural focus in the other. Fifty percent isointensity contours are indicated. **(D)** Multi-slice contrast-enhanced MRI after a sonication that consisted of four different holograms created to match the anatomy in the rat brain. The colored contours indicate the 50% isointensity for the four individual holograms; the white contour is the 50% isointensity contour of the sum of the four sonications. Additional disruption was observed near the skull (*), presumably due to reflections.

Finally, we tested this approach in a macaque. We used a goal in the shape of an ellipsoid with dimensions 3×3×7 mm rotated 20° about the left/right axis Fig. 4A. The volume of the resulting 50% isointensity contour for this focus was 8.1 times larger than that of the natural focus. We sonicated 640 targets in one hemisphere, using the phased array to sequentially target 32 locations per sonication (Fig. 4B). The resulting BBB disruption is evident in the maps of signal enhancement after MRI contrast administration (Fig. 4C). The signal enhancement varied for different brain structures; examples in basal ganglia structures are shown in Fig. 4D-E. The biggest difference was between white and gray matter structures. For example, signal enhancement after contrast injection relative to the non-sonicated hemisphere (39.6 ± 12.7%; p<0.0001) was higher in the caudate than the internal capsule (6.6 ± 4.4%; p<0.0001). The strength of the harmonic emissions, which were used to control the exposure level during sonication, were approximately 5 dB higher than the natural focus (Fig. 4F). No tissue damage was evident in T2*-weighted MRI after sonication (Supplemental Figure 1).

**Fig. 4.**
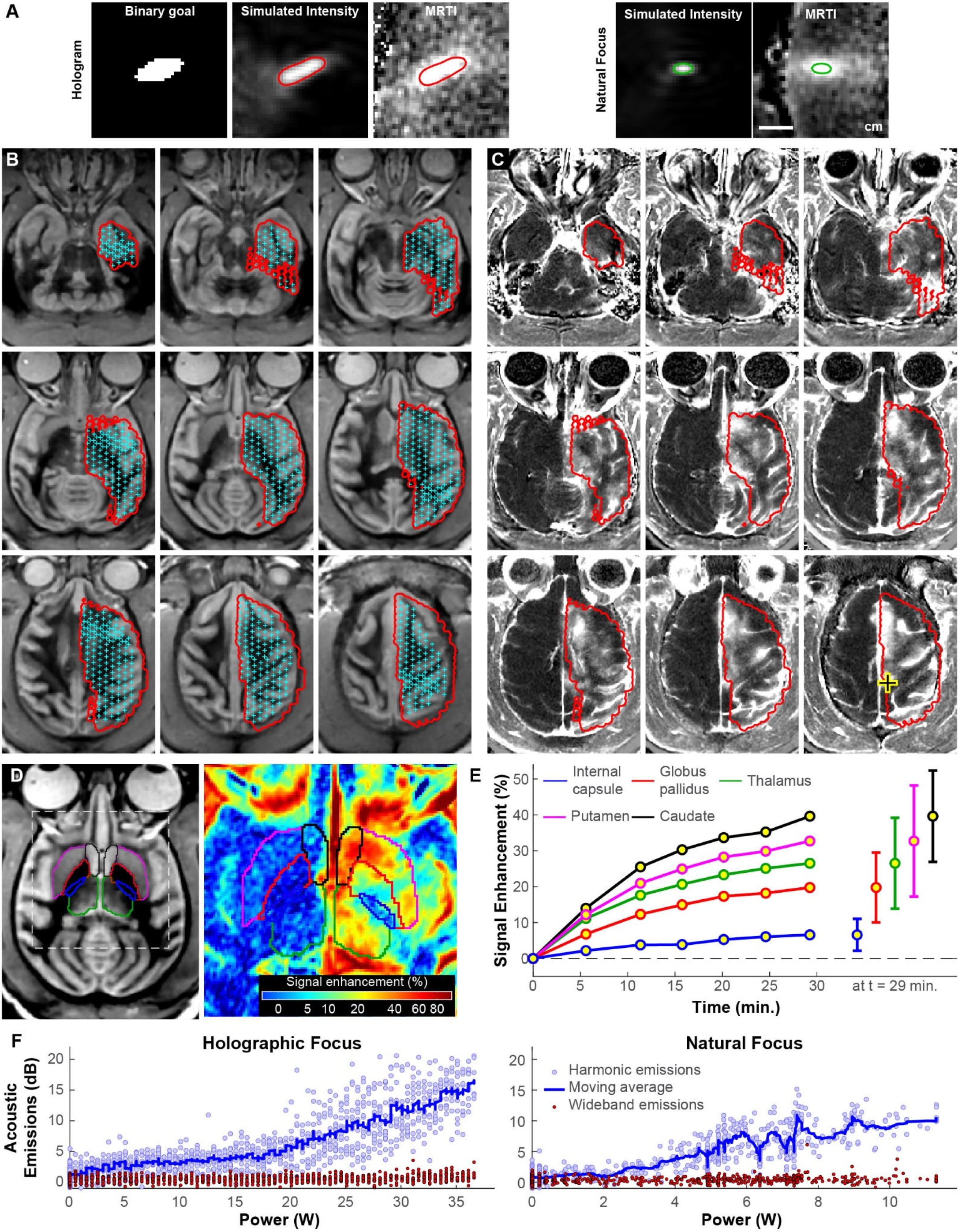
Holographic focusing to disrupt the BBB in one hemisphere in a macaque. **(A)** Sagittal maps (*yz* plane of the transducer) of the simulated intensity (left) and MR temperature imaging (right) of the hologram used in the experiment, which was an ellipsoid rotated 20° about the *x* axis; the natural focus is shown as a reference. **(B)** Axial white matter nulled MRI showing the targets (crosses) that were within 2 mm of each plane; 640 targets were sonicated overall. Fifty percent isointensity contours are shown in red. **(C)** Resulting contrast enhancement after BBB disruption. The location of the geometric focus of the hemispherical transducer is indicated by the cross. **(D)** White matter nulled MRI and enhancement map with different structures segmented in the basal ganglion. **(E)** The enhancement level varied in different structures, with the least disruption in the internal capsule. Enhancement values shown are relative to the non-sonicated hemisphere. **(F)** Acoustic emissions signal strength was increased with enlarged focusing (left) compared to the natural focus (right). Emissions are plotted relative to acquisitions obtained without sonication.

## Discussion

Being able to shape the focal region with a phased array could be useful for patient-specific tailoring of the ultrasound exposures for different brain applications, including BBB disruption, neuromodulation, and ablation. Others have explored using 3D printed acoustic lenses to achieve this result (*3, 13-16*). A phased array has the advantage of being able to generate a different pattern for each sonication. One could design a treatment plan where the focal volume shapes are matched to different brain structures with different vascular densities. Due to such differences, the resulting BBB disruption varies for different structures. Such effects were observed in the monkey experiment, where the disruption varied for different structures despite using the same control settings (Fig. 4D). Improved homogeneity might be achieved by restricting the focal volume to specific brain structures.

Increasing the focal volume could improve our ability to detect small acoustic signals emitted by microbubbles in the brain. More work, however, is needed to use the strength of the acoustic emissions to control the exposure levels for different holograms. One will need to account for the overall volume of the focal region, the relative vascular density of the tissue, and the attenuation of the skull on the emissions signals. The impact of reflections from the inner surface of the skull should also be taken into account if locations near the brain surface are targeted. We observed this effect in the rat experiments.

Including the skull in the procedure and varying the magnitudes of the elements in addition to the phases might be useful in minimizing reflection effects. The element-wise simulation approach used here can include the skull (*17*), and the holography approach could vary the elements’ magnitudes to minimize reflections.

Different methods have been described to create tailored ultrasound field patterns using a phased array transducer (*15, 18-21*). The simple, unguided search scheme described here was developed *ad hoc* by trial and error and produced repeatable results in a few minutes on a desktop PC. Further optimization might yield improved agreement between the goal and the pressure field or faster convergence to a solution. For example, using established optimization methods instead of a blind search could be faster. We also did not simulate the entire acoustic field of the transducer, and there could be hotspots in areas outside of our simulated volume. If that is an issue, repeating the procedure with a larger region of interest should eliminate such hotspots.

Not all three-dimensional pressure distributions are physically possible for a given array, and the method applied here cannot predict beforehand whether the algorithm will converge at an acceptable level. In addition, the maximal size of the focal region is limited by the maximum exposure level that the ultrasound system can produce. With increased volumes, one will also need to take care to monitor the exposure levels in the acoustic beam path to avoid excessive exposures in bone and in non-targeted tissue structures.

In conclusion, a simple holography method was tested with a 1024-element phased array transducer used clinically for BBB disruption. Experiments with MR temperature imaging and BBB disruption in rats and a macaque demonstrated the ability of the approach to create arbitrary focal patterns. We expect that using this approach to shape the focal volume to match different brain structures can reduce the number of sonications needed to disrupt large volumes, improve the homogeneity of the disruption, and improve the sensitivity and specificity of monitoring acoustic emissions to monitor and control the procedure. The approach could also be useful for other ultrasound applications.

## Materials and Methods

### Focused ultrasound system

We used the ExAblate Neuro Type II (InSightec), which uses a 1024-element hemispherical phased array transducer with a diameter of 30 cm and a resonant frequency of 220 kHz. The transducer was not perfectly hemispherical; it consisted of 64 flat tiles each containing nine elements (Fig. 1A). The system was integrated into a 3T MRI (Premiere, GE); imaging was obtained using either a 5×6 cm square or 14 cm diameter circular receive-only surface coil (constructed in-house). For use in preclinical studies, the transducer was placed on its side so that it could be filled with water like a bowl. During the experiments, the system continuously degassed and circulated the water that provided acoustic coupling. The device steered the focal point by adding phase shifts to the array elements so their fields coincided at different locations away from the geometric focus (Fig. 1B). It can also apply phase offsets to correct aberrations induced by the skull bone, but that feature was not used here. The additional phase shifts used to create the holograms were loaded by the device from a text file.

### Acoustic simulations

We modeled the acoustic field using k-Wave, an open-source MATLAB toolbox (22). The acoustic field for one square 1×1 cm flat piston was simulated in water (Fig. 1A). The simulation had a grid size of 76×76×201 with a grid point spacing of 0.97 mm (one-seventh of a wavelength). Density and sound speed in water were 1000 km/m^3^ and 1500 m/s, respectively. We used a perfectly matched layer with an attenuation of 2 Np per grid point and a size of 10 grid points.

We interpolated the complex pressure field for this simulation from the reference frame of each of the 1024 elements of the array (yellow volume in Fig. 1A) to a global 76×76×71 element volume in the *xyz* reference frame of the hemispherical array (green volume in Fig. 1A). The locations of the elements were provided by the manufacturer. The pressure distributions of the 1024 elemental simulations could then be loaded simultaneously into memory. After adding phase offsets to each element, we then preformed a complex sum over the elemental simulations to estimate the combined pressure field (Fig. 1B). We could also modify the magnitude of each element using this approach; such modifications were not employed here.

### Holography

Holograms were created by loading 1024 individual acoustic simulations for the elements of phased array simultaneously into computer memory and systematically modulating the elements’ phases until the combined field matched a desired pattern. We first created a three-dimensional binary goal to locally maximize the pressure amplitude (Fig. 1B). The initial phases for the elements were randomized between zero and 2π. Then, in random order, we selected an element and changed the phase from 0 to 2π in 5 steps. For each step, we summed the 1024 elements to calculate the pressure amplitude in a small region of interest centered on the geometric focus and calculated the cost function described below. If the cost function was reduced, we kept that phase for that element. This procedure, with a completely randomized phase as a starting point, was repeated five times using a small cubic region of interest with dimensions of 10 simulation grid points (9.7 cm), as shown in Fig. 1C. Using the best solution from these five runs as a starting point, we then cycled through the elements with a larger region of interest with dimensions of 20 simulation grid points and evaluated ten phase values between 0 and 2π. Finally, we added random phases between ±2π/3 radians to the elements and cycled through the elements two more times (Fig. 1D).

The cost function was the weighted average of three terms:

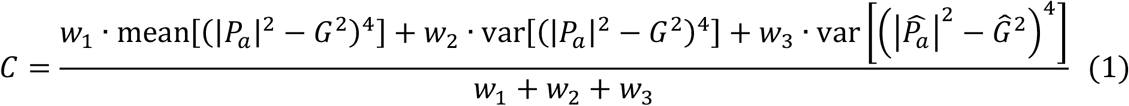

where |*P*_a_| is the magnitude of the normalized simulated pressure field, *G* is the binary goal, and 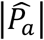 and *Ĝ* are the normalized pressure amplitude and goal within the volume defined by 75% isopressure contour. Weights (*w*_l_, *w*_2_, *w*_3_) were 1, 2, and 10, respectively. This cost function was determined *ad hoc* by trial and error. This procedure was performed in Matlab on a desktop PC (Dell Precision 5820, Intel Core i9-10940X CPU, 3.3 GHz).

### MR temperature imaging

The temperature rise produced during sonications was evaluated in a tissue-mimicking phantom (InSightec) using a gradient echo sequence. Images were obtained in a time series in a single plane every 4.5 seconds. Phase difference images were used to estimate temperature changes (*12*) using a temperature coefficient of -0.0909 ppm/°C. MRI parameters are listed in Supplemental Table 1. Sonication power levels and durations varied depending on the volume of the enlarged focal region; the thermal images served to verify that the holographic focusing was successful.

### Blood-brain barrier disruption

Animal experiments were approved by the institutional review boards of Brigham and Women’s Hospital and Harvard Medical School. Male Sprague Dewey rats were anesthetized using intraperitoneal injections of ketamine (80 ml/kg) and xylazine (10 ml/kg). A male rhesus macaque was anesthetized with intramuscular injection of ketamine (15 mg/kg) and xylazine (0.5 mg/kg). The animal was intubated, and anesthesia was maintained with isoflurane (1.5-2%) and air. The monkey’s heart rate and rectal temperature were continuously monitored throughout the procedure. A heated water blanket was used to maintain body temperature.

Before sonication, the head was shaved, and an intravenous catheter was placed in the tail vein (rats) or leg (monkey). Targets were selected on axial T2*-weighted images. In the monkey, we also acquired a white matter nulled sequence to visualize white and gray matter structures. Sonications (5 ms bursts applied at 1 Hz), were applied to each target in combination with intravenous injections of a microbubble ultrasound contrast agent (Definity, Lantheus). In the rat experiments, we used a bolus injection of 10 μl/kg; they were diluted 10 times. In the macaque studies, 0.8 ml microbubbles were diluted in 4.2 ml saline and preloaded into tubing with a volume of 5 ml that was coiled in a horizontal plane so that they did not migrate after floating. They were administered as an infusion at 12.8 μl/kg/min by pushing them through the tubing using the autoinjector installed at the MRI.

The sonication power level was controlled in real time based on the signal strength of the acoustic emissions obtained at the second and third harmonic of the 220 kHz array. The emissions were captured by an elliptical passive cavitation detector constructed in-house (dimensions: 5×3 cm, resonant frequency: 650 kHz). To avoid vascular damage, power levels were reduced if wideband, subharmonic, or ultraharmonic emissions were detected. Emissions were evaluated in dB relative to measurements obtained without sonication. More details on this controller are described elsewhere (*10*).

In experiments in rats, we compared enlarged focal regions using an elliptical goal with dimensions of 2×3×5 mm. The median power level was 11.0 W [25% quartiles: 10.4 - 12.7 W]; with the natural focus it was 1.5 W [1.1 - 2.0 W]. In addition, we tested holograms with shapes tailored to different anatomical structures (striatum, hippocampus, thalamus, pons), as shown in Fig. 3D. The median power level was 22.4 W [15.1 - 31.2 W].

In the monkey, we tested volumetric opening using an ellipsoid with dimensions of 3×3×7 mm rotated 20° about the left-right axis (Fig. 4A). This angle was similar to the typical angulation of the plane defined by the anterior commissure, posterior commissure, and the midline when the monkey was placed in this setup. We used this hologram to attempt to disrupt the BBB in the entire left hemisphere. We targeted 640 locations selected in axial MRI. The distance between targets was 3 mm in-plane and 4.5 mm between planes. Each sonication targeted 32 targets in sequence. The median power level 44.8 W [25% quartiles: 35.7 - 57.2 W].

T2*-weighted images were acquired after sonication to detect vascular damage. To detect BBB disruption, T1-weighted fast spin echo images were acquired before and after an injection of MRI contrast agent (Gadavist, 0.125 mmol/kg). BBB disruption was evident where the contrast agent extravasated. MRI parameters are listed in Supplemental Table 1.

### Statistical Analysis

Data are presented as mean values ± standard deviation, or median values [25% quantiles]. Enhancement between sonicated and non-sonicated brain regions in the monkey were compared using an unpaired t-test.

## Acknowledgments

The authors thank InSightec for providing the ExAblate Neuro system for preclinical research. In particular, the authors thank Rafi De Picciotto and Javier Grinfeld for their technical help with the system.

## Author contributions

Conceptualization: NM

Methodology: NM

Investigation: NM, ML, SF, YZ

Visualization: NM

Supervision: NM, ML

Writing—original draft: NM

Writing—review & editing: ML, SF

## Data and materials availability

All data, code, and materials used in the analyses that are not proprietary to InSightec are available upon request.

## Supplementary Materials

**Supplemental Table 1:**
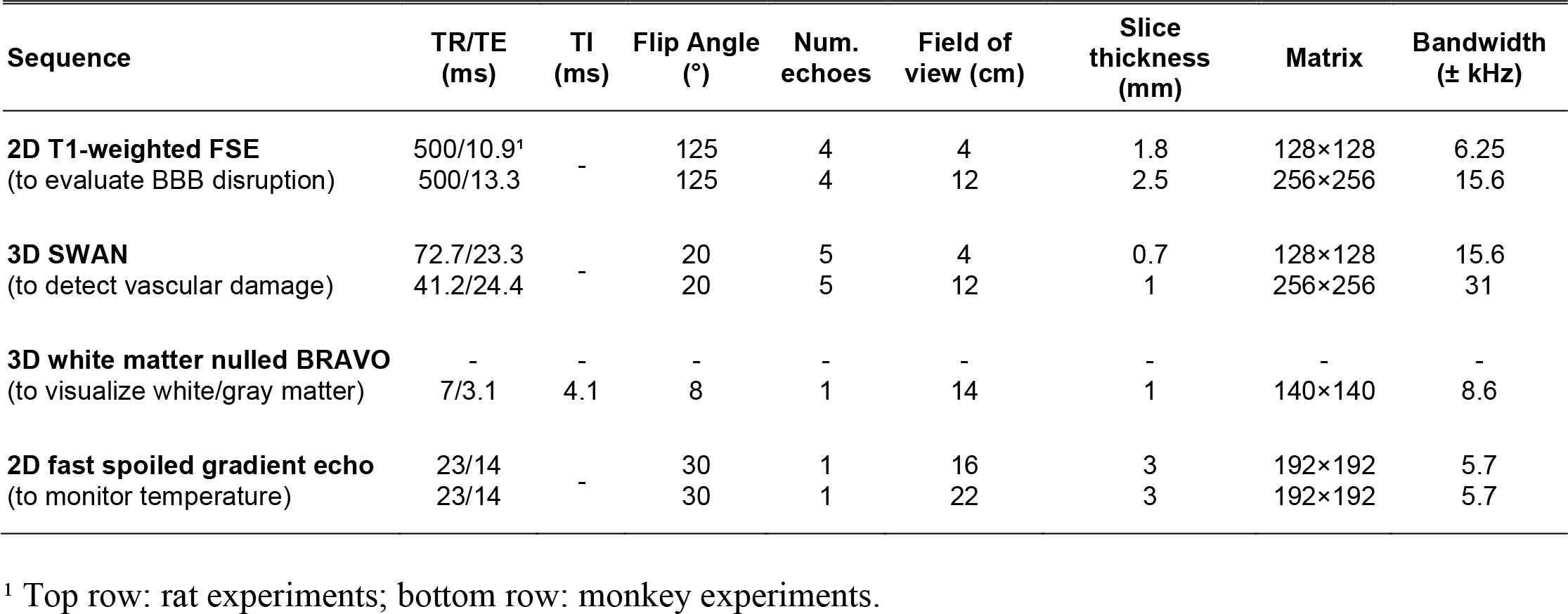
MRI parameters.

**Supplemental Figure 1.**
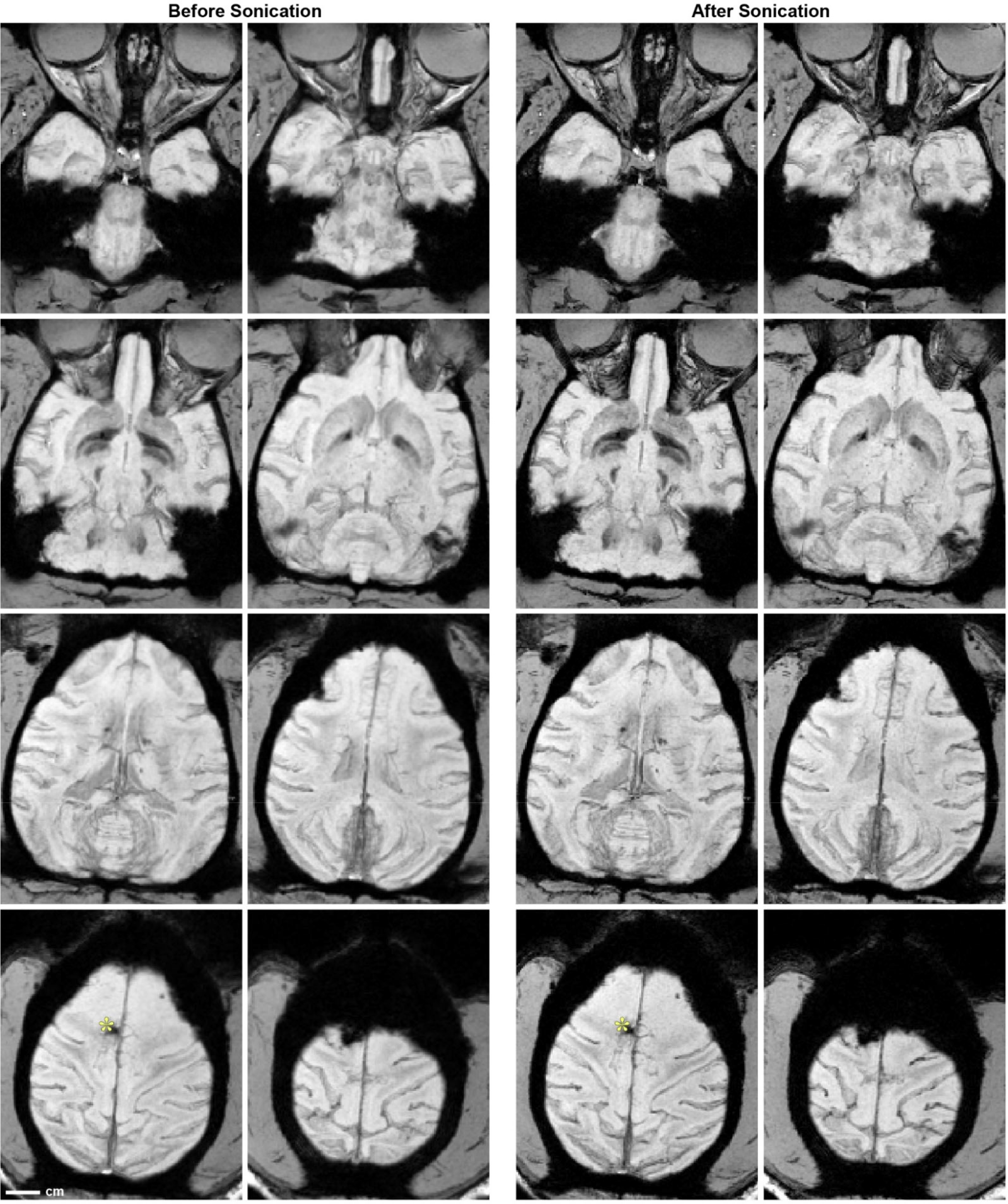
No vessel damage evident in T2*-weighted MRI after sonication in a macaque. Minimum intensity projections over every five images are shown. The monkey had pre-existing brain damage from a prior surgery (*).

## References

1. K. Melde, A. G. Mark, T. Qiu, P. Fischer, Holograms for acoustics. Nature 537, 518–522 (2016).

2. Y. Xie, C. Shen, W. Wang, J. Li, D. Suo, B.-I. Popa, Y. Jing, S. A. Cummer, Acoustic holographic rendering with two-dimensional metamaterial-based passive phased array. Scientific reports 6, 35437 (2016).

3. D. Andrés, N. Jiménez, J. M. Benlloch, F. Camarena, Numerical Study of Acoustic Holograms for Deep-Brain Targeting through the Temporal Bone Window. Ultrasound Med Biol 48, 872–886 (2022).

4. A. Marzo, S. A. Seah, B. W. Drinkwater, D. R. Sahoo, B. Long, S. Subramanian, Holographic acoustic elements for manipulation of levitated objects. Nat Commun 6, 8661 (2015).

5. K. Hynynen, N. McDannold, N. Vykhodtseva, F. A. Jolesz, Noninvasive MR imaging-guided focal opening of the blood-brain barrier in rabbits. Radiology 220, 640–646 (2001).

6. T. Mainprize, N. Lipsman, Y. Huang, Y. Meng, A. Bethune, S. Ironside, C. Heyn, R. Alkins, M. Trudeau, A. Sahgal, J. Perry, K. Hynynen, Blood-Brain Barrier Opening in Primary Brain Tumors with Non-invasive MR-Guided Focused Ultrasound: A Clinical Safety and Feasibility Study. Sci. Rep 9, 321 (2019).

7. N. Lipsman, Y. Meng, A. J. Bethune, Y. Huang, B. Lam, M. Masellis, N. Herrmann, C. Heyn, I. Aubert, A. Boutet, G. S. Smith, K. Hynynen, S. E. Black, Blood-brain barrier opening in Alzheimer’s disease using MR-guided focused ultrasound. Nat Commun 9, 2336 (2018).

8. C. Gasca-Salas, B. Fernández-Rodríguez, J. A. Pineda-Pardo, R. Rodríguez-Rojas, I. Obeso, F. Hernández-Fernández, M. Del Álamo, D. Mata, P. Guida, C. Ordás-Bandera, J. I. Montero-Roblas, R. Martínez-Fernández, G. Foffani, I. Rachmilevitch, J. A. Obeso, Blood-brain barrier opening with focused ultrasound in Parkinson’s disease dementia. Nat Commun 12, 779 (2021).

9. P. Anastasiadis, D. Gandhi, Y. Guo, A. K. Ahmed, S. M. Bentzen, C. Arvanitis, G. F. Woodworth, Localized blood-brain barrier opening in infiltrating gliomas with MRI-guided acoustic emissions-controlled focused ultrasound. Proceedings of the National Academy of Sciences of the United States of America 118, (2021).

10. N. McDannold, Y. Zhang, J. G. Supko, C. Power, T. Sun, C. Peng, N. Vykhodtseva, A. J. Golby, D. A. Reardon, Acoustic feedback enables safe and reliable carboplatin delivery across the blood-brain barrier with a clinical focused ultrasound system and improves survival in a rat glioma model. Theranostics 149, 6284–6299 (2019).

11. N. McDannold, M. Livingstone, C. B. Top, J. Sutton, N. Todd, N. Vykhodtseva, Preclinical evaluation of a low-frequency transcranial MRI-guided focused ultrasound system in a primate model. Phys. Med. Biol 61, 7664–7687 (2016).

12. Y. Ishihara, A. Calderon, H. Watanabe, K. Okamoto, Y. Suzuki, K. Kuroda, A precise and fast temperature mapping using water proton chemical shift. Magn. Reson. Med 34, 814–823 (1995).

13. S. Jimenez-Gambin, N. Jimenez, A. Pouliopoulos, J. M. Benlloch, E. Konofagou, F. Camarena, Acoustic Holograms for Bilateral Blood-Brain Barrier Opening in a Mouse Model. IEEE transactions on bio-medical engineering 69, 1359–1368 (2022).

14. S. Jiménez-Gambín, N. Jiménez, J. M. Benlloch, F. Camarena, Holograms to focus arbitrary ultrasonic fields through the skull. Physical Review Applied 12, 014016 (2019).

15. Y. Hertzberg, O. Naor, A. Volovick, S. Shoham, Towards multifocal ultrasonic neural stimulation: pattern generation algorithms. J. Neural Eng 7, 056002 (2010).

16. G. Kook, Y. Jo, C. Oh, X. Liang, J. Kim, S. M. Lee, S. Kim, J. W. Choi, H. J. Lee, Multifocal skull-compensated transcranial focused ultrasound system for neuromodulation applications based on acoustic holography. Microsystems & nanoengineering 9, 45 (2023).

17. N. McDannold, P. J. White, R. Cosgrove, Elementwise approach for simulating transcranial MRI-guided focused ultrasound thermal ablation. Appl.Phys. Res. 1, 033205 (2019).

18. E. S. Ebbini, C. A. Cain, Multiple-focus ultrasound phased-array pattern synthesis: optimal driving-signal distributions for hyperthermia. IEEE Trans. Ultrason. Ferroelectr. Freq. Contr 36, 540 (1989).

19. S. Elizondo, I. Ezcurdia, J. Goñi, M. Galar, A. Marzo, Enhancing the quality of amplitude patterns using time-multiplexed virtual acoustic fields. Applied Physics Letters 123, (2023).

20. D. Gourevich, Y. Hertzberg, A. Volovick, Y. Shafran, G. Navon, S. Cochran, A. Melzer, Ultrasound-mediated targeted drug delivery generated by multifocal beam patterns: an in vitro study. Ultrasound Med Biol 39, 507–514 (2013).

21. M. H. Lee, H. M. Lew, S. Youn, T. Kim, J. Y. Hwang, Deep Learning-Based Framework for Fast and Accurate Acoustic Hologram Generation. IEEE transactions on ultrasonics, ferroelectrics, and frequency control 69, 3353–3366 (2022).

22. B. E. Treeby, B. T. Cox, k-Wave: MATLAB toolbox for the simulation and reconstruction of photoacoustic wave fields. J. Biomed. Opt 15, 021314 (2010).

